# DataRemix: a universal data transformation for optimal inference from gene expression datasets

**DOI:** 10.1101/357467

**Authors:** Weiguang Mao, Javad Rahimikollu, Ryan Hausler, Bernard Ng, Sara Mostafavi, Maria Chikina

## Abstract

RNAseq technology provides unprecedented power in the assessment of the transcription abundance and can be used to perform a variety of downstream tasks such as inference of gene-correlation network and eQTL discovery. However, raw gene expression values have to be normalized for nuisance biological variation and technical covariates, and different normalization strategies can lead to dramatically different results in the downstream study. We describe a generalization of SVD-based reconstruction for which the common techniques of whitening, rank-*k* approximation, and removing the top *k* principle components are special cases. Our simple three-parameter transformation, DataRemix, can be tuned to reweight the contribution of hidden factors and reveal otherwise hidden biological signals. In particular, we demonstrate that the method can effectively prioritize biological signals over noise without leveraging external dataset-specific knowledge, and can outperform normalization methods that make explicit use of known technical factors. We also show that DataRemix can be efficiently optimized via Thompson Sampling approach, which makes it feasible for computationally expensive objectives such as eQTL analysis. Finally, we apply our method to the ROSMAP dataset and we report what to our knwoledge is the first replicable trans-eQTL effect in human brain.

## Introduction

Genome-wide gene expression studies have become a staple of large-scale systems biology and clinical projects. However, while gene expression is the most prevalent high-throughput technology, technical challenges remain. Raw gene expressio nvalues must be normalized for any technical and nuisance biological variation and the normalization strategy can have dramatic effects on the results of downstream analysis. This is especially true in cases where the sought-after gene expression effects are likely to be small in magnitude, such as expression quantitative trait loci (eQTLs). Increasingly sophisticated normalization methods have been proposed and many are computational intensive and/or can have multiple free parameters that must be optimized (Leek & Storey 2007; Stegle *et al.*. 2010; Listgarten *et al.*. 2010; Kang *et al.*. 2008; Mostafavi *et al.*. 2013). Moreover, it is not uncommon for one dataset to yield multiple normalized versions that maximize performance in a particular setting (such as the discovery of *cis*- and *trans*-eQTLs Battle *et al.*. 2014), highlighting the complexity of the normalization problem.

Singular value decomposition (SVD) is one of the most widely used gene expression analysis tools (Alter *et al.*. 2000, 2003) that can also be used for data normalization. Using the SVD we can simply remove the first few principle components that are presumed to represent technical factors such as batch-effects or other nuisance variation. In some cases this dramatically improves downstream performance, for example in the case of eQTL analysis (Mostafavi *et al.*. 2013). The drawback of this method is that the exact number of components to remove must be determined empirically and some meaningful biological signals may be lost in the process.

More sophisticated approaches attempt to partition data structure into true biological and nuisance variation and remove only the latter (Leek & Storey 2007; Stegle *et al.*. 2010; Listgarten *et al.*. 2010; Kang *et al.*. 2008; Mostafavi *et al.*. 2013). These can improve on the naive SVD-based normalization but require additional input such as technical covariates, or the study design. The success of these methods ultimately depends on the availability and quality of such meta data and some methods still rely on parameter optimization to maximize performance. These widely used normalization approaches all have a common theme that they rely in part on the intrinsic data structure. One key property that contributes to the success of these approaches is that for many biological questions of interest, nuisance variation (of technical or biological origin) is larger in magnitude than true biological variation. Our proposed method, DataRemix, explicitly formalizes this view of the data normalization problem.

In this work we demonstrate that biological utility of gene expression datasets can be dramatically improved with a simple three-parameter transformation, DataRemix. Our method does not require any dataset specific knowledge but rather optimizes the transformation with respect to some independent *objective* of data quality, such as the quality of the gene-correlation network or the number of *trans*-eQTL discoveries. Because our method requires only the gene expression data and biological validity objective, it can be applied to any publicly available dataset. We focus our study on gene expression data for which methods for quantifying biological validity are well established, but our approach can be readily applied to any high-throughput molecular data for which similar quality metrics can be defined. We show that this strategy can outperform methods that make explicit use of dataset specific factors, and can further improve datasets that have been extensively normalized via an optimized, parameter-rich model. We also show how the optimal parameters of DataRemix can be found efficiently by Thompson Sampling with a dual learning setup, making the approach feasible for computationally expensive objectives such as eQTL analysis.

## Result

### The DataRemix framework

We formulate DataRemix as a simple parametrized version of SVD which can be directly optimized to improve the biological utility of gene expression data. Given a gene-by-sample matrix *X*, SVD decomposition can be thought of as a solution to the low-rank matrix approximations problem defined as:

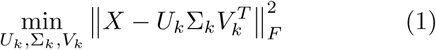

where *U* and *V* are unitary matrices. With the SVD decomposition *U* Σ*V* ^*T*^, the product of *k*-truncated matricies 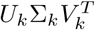 gives the rank-*k* reconstruction of *X*. We introduce two additional parameters *p* and *µ* to define a new reconstruction:

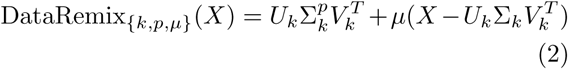

Here, *k* is the number of principle components of SVD and *p* ∈ [−1, 1] is a real number which alters the scaling of each singular value. For *p* = 1, this approach reduces to the original SVD-based reconstruction. For *p* = 0, the transformation gives the frequently used whitening operation (Friedman 1987). As depicted in Figure 1, generally, different choices of *p* reweigh the contribution of each variance component, possibly making some low-variance biological signals visible while down-weighting technical and other systematic noise. The parameter *µ* is a non-negative weight that adds the residual back to the reconstruction in order to make the transformation *lossless*.

**Figure 1:**
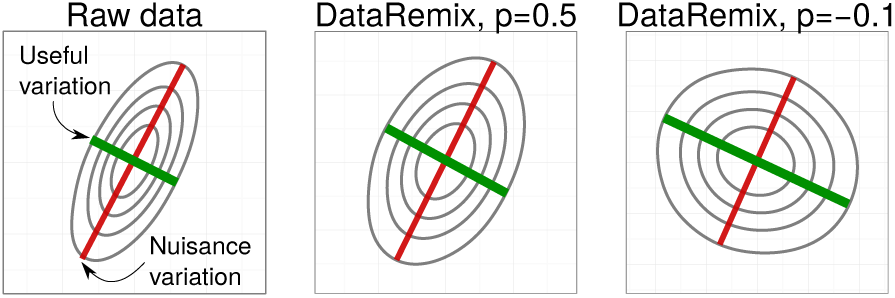
Visual representation of DataRemix transformation. We simulate a 2-dimensional dataset where the nuisance variation contributes more variance than true biological variation. Different power parameters *p* reweigh the contributions of the two variance axes, making the true biological variation more “visible”.

Intuitively, we expect this approach to succeed because sophisticated normalization methods that use both data structure and some external variables, such as technical covariates, can be thought of as implicit regularizations on the naive SVD-based normalization (which simply removes the first *k* components), and this formulation simply makes this explicit.

The general workflow of DataRemix is shown in Figure 2. The downstream biological objective depends on your study. For example, if you focus on *trans*-eQTL analysis, the biological objective will be to increase the number of *trans*-eQTLs detected from the DataRemix-normalized gene expression profile and the metric *y* will be the number of *trans*-eQTLs deemed significant. The parameter optimization step which determines the next point to check is detailed in the Methods section.

**Figure 2:**
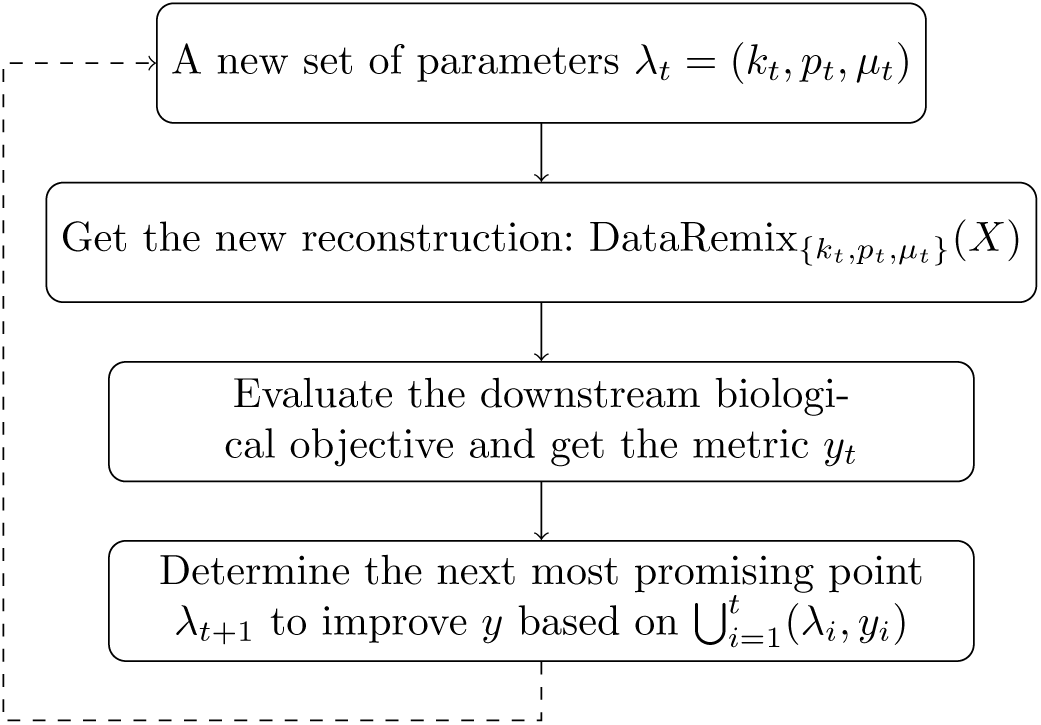
The workflow of DataRemix.

### Quality of the correlation network derived from the GTEx gene expression study

The GTEx datasets (Lonsdale *et al.*. 2013) is comprised of human samples from diverse tissues, many of which were obtained post-mortem and there are many technical factors which have considerable effects on the gene expression measurements. On the other hand this rich dataset provides an unprecedented multi-tissue map of gene regulatory networks and has been extensively analyzed in this context. It is natural to assume that a dataset that is better at recovering known pathways is likely to yield more credible novel predictions. Thus, we use DataRemix to optimize the known pathway recovery task as a function of the correlation network computed on a Remixed dataset.

We formally define the objective as the average AUC across “canonical” mSigDB pathways (which include KEGG, Reactome and PID) (Subramanian *et al.*. 2005) using guilt-by-association. Specifically, the genes are ranked by their average Pearson correlations to other genes in the pathway (excluding the gene when the gene itself is a pathway member). Figure 3A depicts the results of grid search for the parameters *k* and *p* (with *µ* fixed at 0.01) and the contour plot shows a clear region of increased performance. Using the optimal transformation found by grid search, we plot per-pathway AUC improvement in Figure 3B and find that the AUC is substantially increased for almost every pathway.

**Figure 3:**
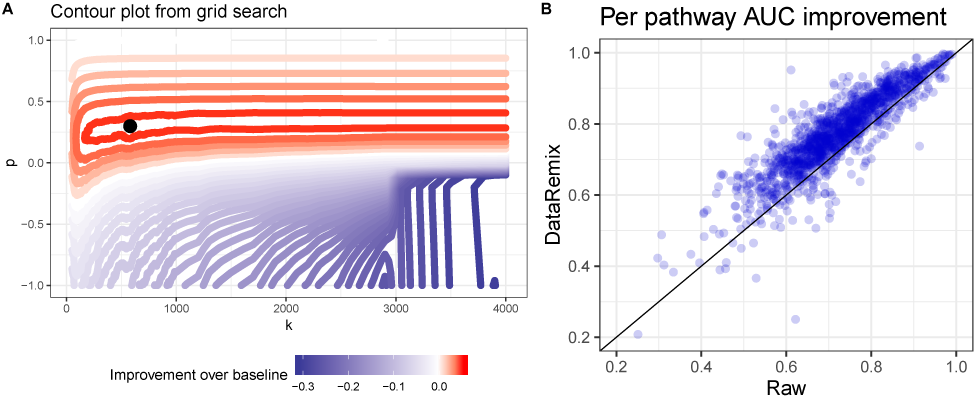
**A** The improvement in performance of DataRemix transform of the pathway prediction task visualized as a function of *k* and *p* parameters (*µ* is fixed at 0.01). Performance is measured as the mean AUC across all pathways in the “canonical” mSigDB dataset and the red contours indicate improvement over the performance on untransformed data. **B** Per-pathway performance improvement for the DataRemix transformation corresponding to the optimal point in **A**.

In Figure 4 we systematically evaluate the performance of DataRemix against alternative methods. For the purpose of evaluation we include the naive method of simply removing known and hidden factors from the data. We consider removing principle componentes (Remove PC), removing known technical variables (Remove tech), and a combination of the two (Remove tech and PC). Since the number of hidden factors is not known, we optimize the number of PCs removed to the specific network quality objective (see Methods for further details). We also include a penalized mixed linear model method “Hidden Covariates with Prior” (HCP) which takes known covariates as input. In addition to the number of hidden components, this method has 3 hyper-parameters that were optimized to maximize the network quality objective via grid search. HCP has been extensively benchmarked perviously and has been shown to outperform both naive methods and the widely used PEER approach (see (Stegle *et al.*. 2010) for PEER and (Mostafavi *et al.*. 2013) for HCP including performance comparison). Moreover, HCP is considerably faster than PEER making an extensive hyper-parameter search feasible.

**Figure 4:**
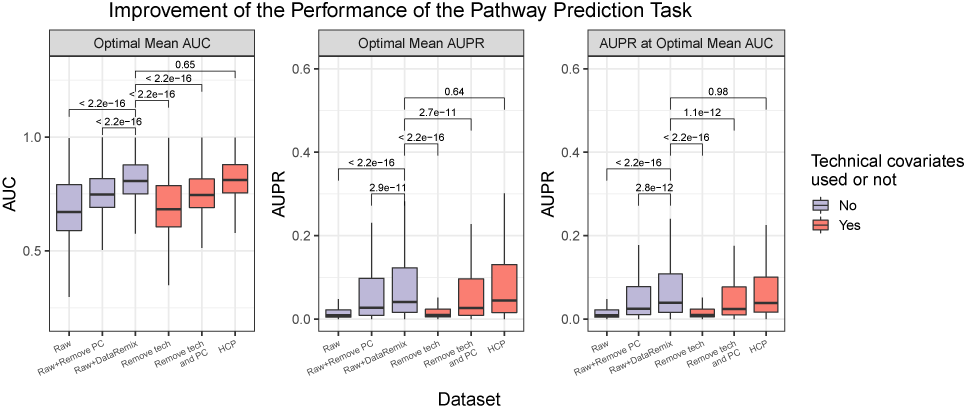
We compare our DataRemix approach to other common normalization strategies with respect to correlation network quality. Here, we consider different normalizations of the GTEx dataset and the details are described in Table. 2. We compute several “naive” normalizations which simply remove known factors (tech), top *k* principle components where *k* is optimized for the task (PC) or both (tech+PC). We also consider “Hidden Covariates with Prior” (HCP) which is a mixed linear model that takes known factors into account and has been shown to outperform other methods in various normalization tasks (Mostafavi *et al.*. 2013). The four hyper-parameters in HCP are optimized by grid search. Each box plot shows the distribution in AUCs or AUPRs across the “canonical” mSigDB pathways. P-values compare the results achieved by DataRemix against others using the Wilcoxon ranksum test. DataRemix’s performance surpasses all naive methods and is comparable to HCP while using no technical covariates and considerably less computation (see text for details).

We find that on this dataset DataRemix is able to outperform all naive methods including ones that make use of known technical covariates, achieving performance that is comparable to that of HCP. In summary, our DataRemix framework is able to match the performance of the best competing method, HCP, *while using no technical covariates*. It is worth pointing out that once a truncated SVD decomposition is computed, a single DataRemix evaluation requires only two matrix multiplications while HCP is an optimization problem which needs to be solved iteratively with two matrix inversions at each step.

### eQTL discovery in the DGN dataset

We also consider the task of discovering *cis*- and *trans*-eQTLs on the Depression Gene Networks (DGN) dataset (Battle *et al.*. 2014). In the original analysis this dataset was normalized using the Hidden Covariates with Prior (HCP) (Mostafavi *et al.*. 2013) with four free parameters that were separately optimized for *cis*- and *trans*-eQTLs. The rationale behind separate *cis* and *trans* optimized normalization can be understood in terms of which variance components represent true biological vs. nuisance variation in the two contexts. Specifically, *cis*-eQTLs represent *direct* effects of genetic variation on the expression of a single gene. On the other hand, *trans*-eQTLs represent network level, *indirect* effects that are mediated by a regulator. Thus, *trans*-eQTLs are reflected in systematic variation in the data which becomes a nuisance factor when only direct effects are of interest. It thus follows that the data should be more aggressively normalized for *cis*-eQTL discovery. The original analysis of this dataset optimized the HCP parameters separately for the *cis* and *trans* tasks yielding two different datasets that we refer to as *D*_HCP−cis_ and *D*_HCP−trans_.

The HCP model takes various technical covariates as input, and 20 of the covariates used in the original study cannot be inferred from the gene-level counts. In order to investigate how much improvement can be achieved via DataRemix in the absence of access to these covariates, we also consider a “naively” normalized dataset, quantile normalization of log-transformed counts, or *D*_QN_.

#### cis-eQTLs

In this task we focus on optimizing the discovery of *cis*-eQTLs. We define *cis*-eQTLs as a SNP-gene interaction where the SNP is located within 50kb of the gene’s transcription start site. The interaction is quantified with Spearman rank correlation and deemed significant at 10% FDR (Benjamini-Hochberg correction for the total number of tests).

We perform our analysis in a cross-validation framework, whereby we optimize DataRemix parameters (using grid search or Thompson Sampling) using SNPs on the odd chromosomes and then evaluate the parameters on the, held-out, even chromosome set. Since there are no hyper-parameters to optimize the even chromosome validation is performed exactly once.

The final results for both the train and test set are depicted in Figure 5. As expected, the quantile-normalized dataset *D*_QN_ performs considerably worse than *D*_HCP−cis_, which is specifically optimized for *cis*-eQTL detection. However, the two datasets achieve comparable performance after applying DataRemix. Moreover, the final performance of the Remixed *D*_QN_ dataset is an improvement on *D*_HCP−cis_ demonstrating the near optimal normalization is possible without access to technical covariates. Importantly, we find that the optimal parameters are indeed generalizable as we achieve a similar level of improvement on the train and test chromosomes.

**Figure 5:**
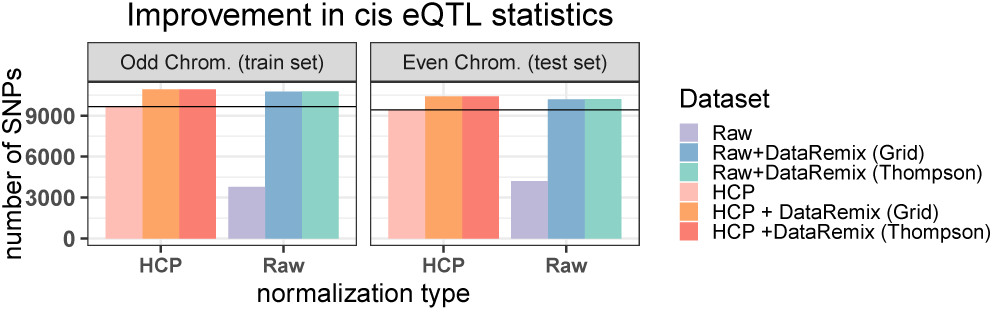
Final results from DataRemix parameter search using a cross-validation framework. *cis*-eQTL statistic is defined to be number of SNP-gene interaction deemed significant at 10% FDR (Benjamini-Hochberg correction for the total number of tests), where the SNP is located within 50kb of the gene’s transcription start site. Optimal parameters are determined using the odd chromosome SNPs only and then tested on the even chromosome SNPs. While the raw dataset is considerably worse than HCP, both are improved to a similar level with DataRemix. We find that the DataRemix transform does not overfit the objective as the degree of improvement is similar across the test and train SNP sets. (Note, the starting value of the raw or HCP dataset differ between the test and train SNP set). Moreover, we find that Thompson Sampling is able to match grid search results using only 100 evaluations.

#### trans-eQTLs

In our second task, we optimize the discovery of *trans*-eQTLs in the same DGN dataset. Ideally, *trans*-eQTLs represent network-level effects and thus give some insight about the regulatory structure of gene expression. However, in practice *trans*-eQTLs are simply defined as SNP-gene associations where the SNP and the gene are located on different chromosomes. While this is a useful heuristic definition, it doesn’t guarantee that the association is mediated at the network level. One possible source of bias is mis-mapped RNAseq reads which contaminate the quantification of the apparently *trans*-associated gene with reads from a homologous locus that has *cis* association. Even in the absence of technical artifacts, direct interchromsomal interactions have been observed (see Williams *et al.*. 2010 for a comprehensive review). In order to focus on potential indirect effects, we apply an additional filter to *trans*-eQTL discovery. Specifically we require SNPs involved in a *trans* effect to be associated with more than one gene at a FDR of 20% (Benjamini-Hochberg correction for the total number of tests (approximately 8 × 10^9^). We term these SNPs *trans*-SNPs^+^. In comparison with same chromosome *cis*-eQTLs, inter-chromosome *trans*-eQTLs are rare and *trans*-SNPs^+^ (as defined above) are more rare still. In fact, using the odd chromosome SNPs subsampled at 20%, we find only 88 such SNPs using *D*_HCP−trans_ dataset and this is the default value we wish to improve.

Here again we find that the dataset specifically optimized for the task of *trans*-eQTL detection, *D*_HCP−trans_, considerably outperforms the raw data *D*_QN_, however DataRemix is able to improve both to a similar performance. As is the case with the *cis*-eQTL objective, the cross-validation procedure gives consistent results and no overfitting is observed for either grid search or Thompson Sampling (Figure 6). We note that Thompson Sampling is able to achieve a better performance than grid search, though the improvement is small in absolute magnitude due to the scarcity of *trans*-eQTLs. In this case, the optimal region for the DataRemix transformation is relatively small (Supplementary Figure S3) and thus Thompson Sampling has an advantage since it can search off the grid.

**Figure 6:**
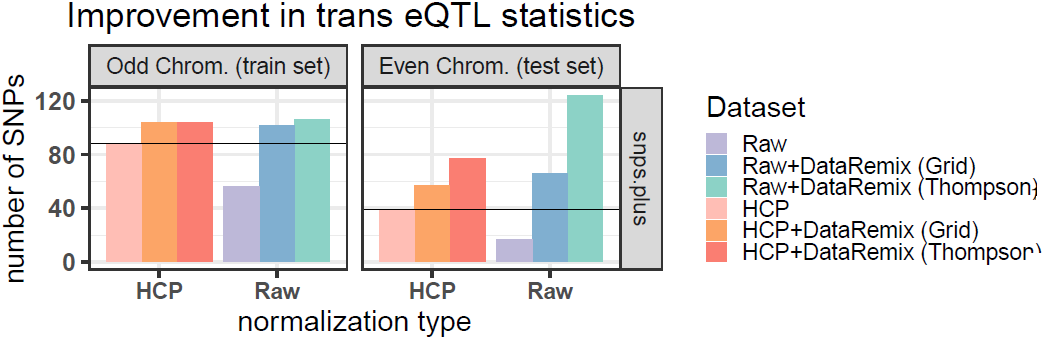
Final values for the eQTL statistics obtained from two versions of datasets. *trans*-eQTL statistic is defined to be number of SNPs involved in a *trans* effect and associated with more than on gene at a FDR of 20% (Benjamini-Hochberg correction for the total number of tests). Here we make a comparison between quantile normalized *D*_QN_ and HCP normalized *D*_HCP−trans_ with parameters optimized for *trans*-eQTL discovery. We find DataRemix is able to improve upon either of starting datasets and the improvements on both the train and test dataset are comparable which indicates that overfitting is not a problem

#### DataRemix performance transfers across different network objectives

It is well know that for statistical analyses of genomic datasets, more significant associations do not necessarily mean improved biological findings. However, it is generally agreed that improvement in *cis*-eQTL detection cannot be achieved through artificial means but indeed represents improved correction for confounding factors (Stegle *et al.*. 2010; Mostafavi *et al.*. 2013). There is no such consensus for *trans*-eQTLs which are rare, and subject to many artifacts. Consequently, it is important to further corroborate the biological validity of the *trans*-optimized dataset through independent means.

Since *trans*-eQTLs are likely to reflect pathway-level effects, we expect that a dataset that is optimally transformed for *trans*-eQTL discovery should also produce better correlation networks. We thus investigate if optimal DataRemix transform is transferable across these tasks by verifying that the Remixed dataset optimized with respect to *trans*-eQTL discovery also improves the network quality criterion. Similar to our analysis of the GTEx datasets, we use the correlation network to perform guilt-by-association pathway predictions and evaluate the results over 1,330 MSigDB canonical pathways. Figure 7 shows scatter plots of per-pathway AUPR (area under precision-recall curve) for several comparisons with respect to the baseline *D*_HCP−trans_ dataset. In the first panel we contrast the performance to *D*_QN_ and observe that, as expected, *D*_HCP−trans_ brings a considerable improvement over the quantile normalized dataset. In the second panel we contrast *D*_HCP−trans_ with the Remixed version of *D*_QN_ (optimized for *trans*-eQTL discovery with Thompson Sampling). We find that the pattern becomes opposite and the Remixed *D*_QN_ dataset performs consistently better that *D*_HCP−trans_. The final panel shows the results of Remixing *D*_HCP−trans_ itself which also improves the performance.

**Figure 7:**
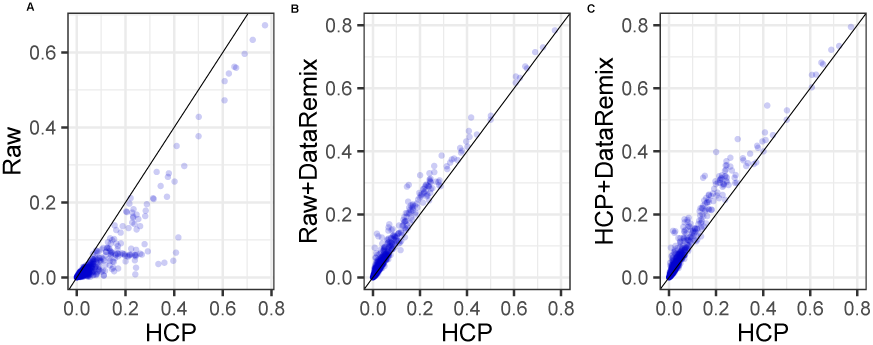
DataRemix-transformed datasets improve the pathway prediction objective which is not explicitly optimized. Each plot is a per-pathway AUPR (area under precision-recall curve) from various datasets (y-axis) contrasted with the results from the optimal covariate-normalized dataset *D*_HCP−trans,_ which serves as the baseline (x-axis). Panel A shows the contrast between *D*_HCP−trans_ and *D*_QN_. The performance of *D*_HCP−trans_ is considerably better. Panel B shows the results of the Remixed *D*_QN_ datasets (optimized for *trans*-eQTL discovery with Thompson Sampling). Even though *D*_QN_ starts out as considerably worse, the Remixed version is able to outperform *D*_HCP−trans_. Panel C shows the results of Remixed *D*_HCP−trans_. We choose to show AUPR instead of AUC because we find that Remixed version matches but doesn’t further improve the AUC performance of *D*_HCP−trans_

Overall, we find that DataRemix improves multiple criteria of biological validity as optimizing for the *trans*-eQTL objective also results in improved correlation networks.

A major finding of our study is that for the eQTL and pathway prediction tasks, the starting point of normalizing DGN datasets appears to matter relatively little. Even though the quantile-normalized dataset performs considerably worse in the beginning, after Remixing its performance matches that of the optimal covariate-normalized datasets. Of course, if covariates are available, it is preferable to use them and in the case of DGN, slightly further improvement can be achieved. However, our results indicate that in some cases datasets *can* be effectively normalized even in the absence of meta-data about quality control or batch variables. This is an important consideration for many legacy datasets where such information is not available.

### Novel Biological Findings

#### New trans-eQTL effects in the DGN dataset

At the optimal DateRemix parameters for *D*_QN_, we find 3000 gene-SNP trans associations at a Benjamini-Hochberg FDR of 0.2 where in contrast to 1691 for *D*_*HCP* −*trans*_. We verified the replication of these associations in an independent dataset, NESDA and find that 1013 (33%) of the DataRemix associations had a replication FDR of < 0.2 while for the default *D*_*HCP* −*trans*_ dataset the same number was 707 (41%). The replication rate was somewhat smaller on the Remixed dataset, which is expected as the replication was performed on raw NESDA data. However, the *total* number of replicated effects was greater.

We highlight an example of new regulatory module recovered via DataRemix that appears to be biologically credible based on independent replication and the known functions of the genes involved. We find that SNP rs11145917 located near CARD9 gene is associated with three genes in the alpha interferon response. The locus has been associated with Crohn’s disease (Franke *et al.*. 2010) and Ulcerative colitis (Anderson *et al.*. 2011) though to our knowledge no mechanism has been proposed. We find that rs11145917 has a cis effect on CARD9 and the trans effects are partially mediated by CARD9 expression. In summary, our analysis suggests that CARD9 may affect baseline activity of the alpha interferon pathway, which is a testable prediction with potential clinical importance.

#### Analysis of the Religious Orders Study and Memory and Aging Project (ROSMAP) Study

We sought to apply our method to the Religious Orders Study and Memory and Aging Project (ROSMAP) Study dataset which consists of 370 human samples with paired gene expression and genotype information. To our knowledge no trans-eQTLs have been reported for human brain and indeed we could not detect any genome-wide significant trans effects in the ROSMAP dataset. Since no trans-eQTLs can be detected, there is no variance in this objective and thus our method cannot be applied directly. However, using the DGN dataset we have shown that optimizing for trans-eQTL discovery also optimizes the network quality objective demonstrating that the two objectives are related. Thus, for the ROSMAP dataset we can optimize network quality (which is quantitative and thus always has some variance across DataRemix parameter settings) and hope to implicitly optimize trans-eQTL discovery. Figure 8 A shows the change in mean AUC and mean AUPR for the network objective after applying DataRemix (see Methods for details). We find that while the mean AUC changes modestly the mean AUPR is nearly doubled. Applying trans-eQTL analysis to the Remixed ROSMAP dataset we detect a single significant effect between CYP2C8 (chr10) and rs10821352 (chr9). This effect was replicated in the CommonMind Consortium dataset (Fromer *et al.*. 2016) with a p-value of 3.1382e-16 (Spearman rank correlation). The interaction passed all quality checks. Specifically, all CYP2C8 30-mers mapped back to CYP2C8 indicating that artifacts from mismapped reads were unlikely and furthermore the eQTL effect was consistent across all 8 exons (Figure S1). To our knowledge this is the first replicated trans-eQTL reported in human brain data.

**Figure 8:**
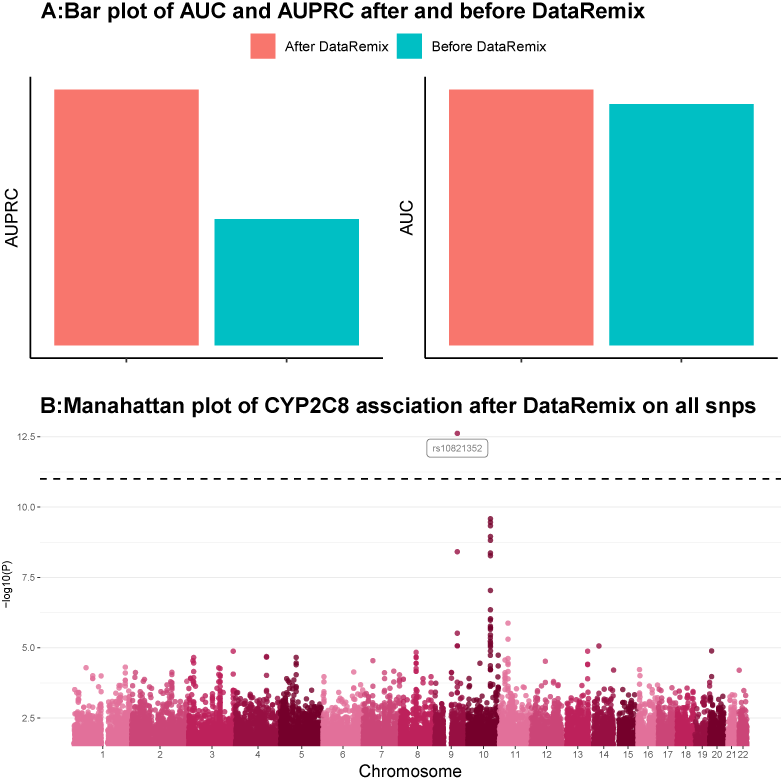
A. Improvement in the network quality objective after running DataRemix with Thompson sampling. B. Manhattan plot of associations with CYP2C8 expression. The CYP2C8 gene is located on chromosome 10. A single SNP on chromosome 9 shows a strong trans effect with a p-value that is notably smaller than the group of cis-effect SNPs on chromosome 10.

The gene, CYP2C8, is a member of the cytochrome P450 and is thought to be involve in the metabolism of polyunsaturated fatty acid and lipophilic xeon-biotics. The xenobiotic metabolism function is supported by the correlation network around CYP2C8. Among its top neighbors is GSTA4 (rank 1, Spearman *ρ* =0.68), CES4A (rank 4, Spearman *ρ* =0.66) two other genes implicated in xenobiotic metabolism. The precise mechanistic nature of how genotype in the rs10821352 locus affects CYP2C8 expression is unclear. No cis-eQTLs for rs10821352 could be detected in ROSMAP and none are reported in the GTex brain data.

### Simulation Study

In order to evaluate the performance of DataRemix when different variance components align with the true biological signals, we performed a simulation study focusing on three representative cases. The cases are: 1) only high-variance components encode biological signals (high-variance Figure 9), 2) only low-variance components encode biological signals (low-variance) and 3) both high- and low-variance components correspond to useful variations (general case). We simulated gene expression profile along with ground-truth pathways and evaluated whether DataRemix could improve the recovery of the simulated pathways (AUC and AUPR) using guilt-by-association.

**Figure 9:**
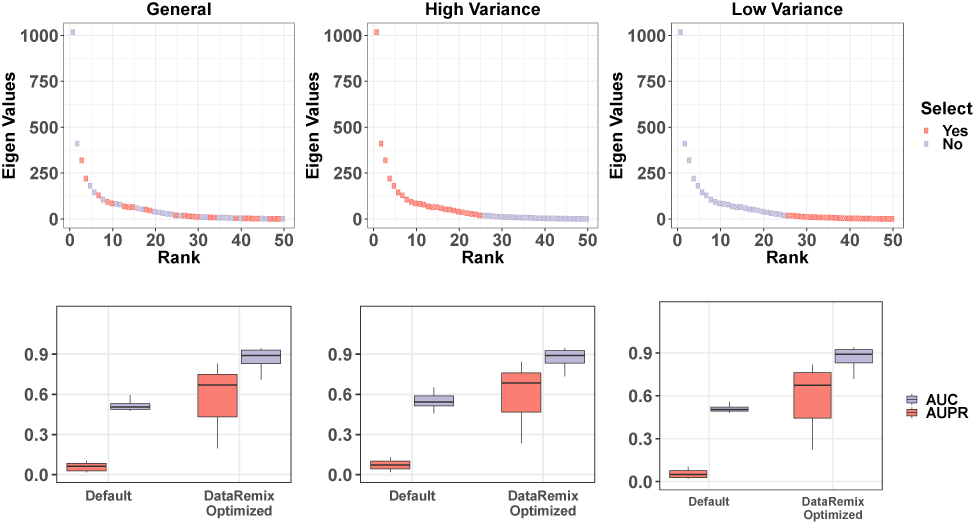
We simulate gene expression data with a low rank approximation so that the component variance distribution approximates that which is typically seen in gene expression data (top row). According to our assumptions only some of the low rank components represent useful biological variation. The left, middle and right panel depict the general, high-variance and low-variance case with the pink points denoting the factors with biological variations. These factors are used to construct the ground truth pathway membership matrix. In the second row, we compare the AUC and AUPR for recovering the pathway co-membership via guilt-by-association analysis on the correlation network. DataRemix is able to improve this metric by reweighing the contribution of different variance components.

**Figure 10:**
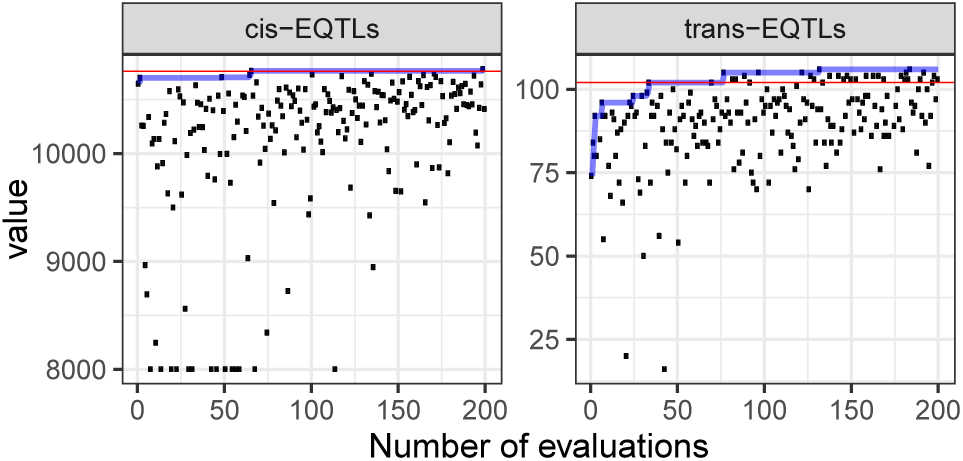
Objective evaluations as a function of iteration number for the *trans*-eQTL and *cis*-eQTL objectives using the quantile normalized *D*_QN_ dataset. Red lines indicate the maximum value that was obtained by grid-search and blue lines indicate the cumulative maximum of Thompson Sampling.

**Table 1:** The association of rs11145917 with genes in the alpha interferon pathway is replicated in an independent dataset. We note that the FDRs for the NESDA dataset represent a correction for the total number of replication test performed, that is only gene-SNP pairs that passed an FDR < 0.2 in the DGN dataset. Since the fraction of true positives in the the replication scenario is higher, the FDRs are lower than the genome wide FDRs at the same p-value.

| SNP | Gene | Method | Spearman rho | p-value | FDR(B.H.) |
| --- | --- | --- | --- | --- | --- |
| rs11145917 | SIGLEC1 | DataRemix | -0.1782 | 5.1052E-08 | 0.0889 |
|  |  | Raw | -0.1510 | 4.1326E-06 | >0.2 |
|  |  | NESDA replication | -0.0414 | 8.1499E-02 | 0.2148 |
|  | IEIT1 | DataRemix | -0.1783 | 5.0403E-08 | 0.0881 |
|  |  | Raw | -0.1627 | 6.7749E-07 | >0.2 |
|  |  | NESDA | -0.07919 | 8.6260E-04 | 0.0050 |
|  | ISG15 | DataRemix | -0.1830 | 2.1867E-08 | 0.0451 |
|  |  | Raw | -0.1541 | 2.5755E-06 | >0.2 |
|  |  | NESDA replication | -0.07451 | 1.7229E-03 | 0.0088 |

We simulated gene expression profile with 5000 genes, 300 samples and 50 latent factors based on the following linear model.

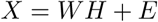

We set *W* and *H* to be positive. Each column of *W* and each row of *H* was drawn from a Normal distribution with mean equal to zero, and the variance parameters were drawn from Exponential distribution with 1e-3 as rate. In this way, the singular values can decrease gradually as the rank increases and each latent factor can have a non-negligible effect when recovering simulated pathways. The matrix *E* ∈ 𝒩 (0, 2) represents random noise.

The gene expression profile is consistent across three cases and a different pathway matrix is generated separately according to each assumption. In the high-variance case, we select the top 25 latent factors. In the low-variance case we pick up the last 25 latent factors and randomly sample 25 latent factors for the general case. Then for corresponding columns in *W*, we randomly select a threshold between 0.01 and 0.1 with 0.01 as the step size. With the threshold value, we pick up the corresponding highest quantile of genes to construct the pseudo geneset as ground truth. The simulated data is used to construct a gene-correlation network which is evaluated according to guilt-by-association recovery of the ground-truth pathways, a commonly accepted network quality metric. We evaluate both the raw data and the optimized Remixed result. In all 3 cases DataRemix was able to substentially improve network quality metrics.

### Thompson Sampling Performance

We find that Thompson Sampling matches the best grid-search performance in under 100 steps giving a 40-fold reduction in the number of evaluations. We also note that it is possible for the Thompson sampling to surpass the grid-search results since the parameter combinations are not constrained by the choice of grid.

## Discussion

We have proposed DataRemix, a new optimizable transformation for gene expression data. The transformation is able to improve the biological validity of gene expression representations and can be used for effective normalization in the absence of any knowledge of technical covariates. One limitation of the DataRemix approach is that it works best on data that is well approximated by a single Gaussian. However, it is relatively straightforward to adapt the approach to matrix decompositions different from SVD that are more suitable for non-Gaussian data, such as independent component analysis. We also note that it is possible to introduce additional parameters that specify more complex weighting schemes. However, as the number of parameters is increased, there is a potential for over-optimization of a specific objective above others. We emphasize that in our simple parametrization, we observe that multiple metrics of biological validity improve when only one is explicitly optimized. Specifically we find that optimizing for *trans*-eQTL discovery also improves the correlation network as measured by guilt-by-association pathway prediction. This property is less likely to be preserved as the number of parameters is increased.

## Methods

### GTEx Dataset

We downloaded the complete gene-level TPM data (RNASeQCv1.1.8) from the GTEx consortium (Lonsdale *et al.*. 2013). These data were quantile normalized to create the raw dataset. We subsequently subjected the dataset to several different normalization approaches that account for hidden and known technical factors.

The technical covariates selected were those with the median values of the variance they explained across genes that were above 0.01. The 8 variables that met this threshold were: SMTS (Tissue type, area from which the tissue sample was taken), SMTSD (Tissue type, more specific detail of tissue type), SMUBRID (Uberon ID), SMNABTCHT (Type of nucleic acid isolation batch), SMEXNCRT (Exonic Rate: the fraction of reads that map within exons), SMGNSDTC (Genes detected), SMTRSCPT (Transcripts detected) and SMNTRNRT (Intronic Rate: the fraction of reads that map within introns).

### DGN Dataset

Depression Gene Networks (DGN) dataset contains whole-blood RNA-seq and genotype data from 922 individuals. The genotype data was filtered for MAF>0.05. The genomic coordinate of each SNP was taken from the Ensembl Variation database (version 90, hg19/GRCh37). SNP identifiers that were not present in that release were excluded. After filtering, there were 649,875 autosomal single nucleotide polymorphisms (SNPs). Data is available upon application through NIMH Center for Collaborative Genomic Studies on Mental Disorders. For gene expression we used the gene-level quantified dataset. The dataset came already filtered for expressed genes and was further filtered for gene symbols that were not present in Ensembl 90 leaving 13,708 genes. The dataset comes in two covariate normalized versions with normalization parameters optimized for *cis-* and *trans*-eQTL discovery separately. To create the naive-normalized dataset, we applied a log transformation, *log*(*x* + 1), to the raw counts and quantile normalized the results.

### ROSMAP dataset

The raw data was obtained from Synpase (syn3219045). The data was optimized for the network quality objective using the canonical pathway genesets from MSigDB (Subramanian *et al.*. 2005). The data was corrected for sex, age and 10 genotype principle components. In order to quantify exon-level effects we used the Synapse BAM files to quantify exon-level FPKMs using featureCounts (Liao *et al.*. 2013).

### NESDA

The NESDA (Netherlands Study of Depression and Anxiety) dataset was obtained from dbGAP (phs000486.v1). Following suggestions from study authors, the NESDA dataset was normalized for sex,age, and the first 10 genotype PCs using linear regression. Genotypes were imputed using Michigan Imputation Server (Das *et al.*. 2016) using 1000 Genome Phase 3 (Version 5) as the reference panel. We assesed the replication of DGN eQTLs based on exact gene and SNP matches.

### Correlation network evaluation

We evaluated the quality of the correlation network derived from a particular dataset using guilt-by-association pathway prediction. Specifically, the genes were ranked by their average Pearson correlations to other genes in the pathway (excluding the gene when the gene itself is a pathway member). The resulting ranking was evaluated for performance using AUC or AUPR metric. For pathway ground-truth, we used the “canonical” pathways dataset from MSigDB, comprising 1,330 pathways (Subramanian *et al.*. 2005).

### eQTL mapping

eQTL association mapping was quantified with Spearman rank correlation. For *cis*-eQTLs, testing was limited to SNPs which locate within 50kb of any of the gene’s transcription start sites (Ensembl, version 90). *cis*-eQTl is deemed significant at 10% FDR with Benjamini-Hochberg correction for the total number of tests. For *trans*-eQTLs, the significance cutoff is 20% FDR with Benjamini-Hochberg correction for the total number of tests. Since the Benjamini-Hochberg FDR is a function of the entire p-value distribution in order to ensure consistency comparisons, the rejection level was set once based on the p-value that corresponded to 10% or 20% FDR in the original *cis*-optimized *D*_HCP−cis_ and *trans*-optimized *D*_HCP−trans_ dataset respectively. To reduce the computational cost of grid evaluations, all the optimization computations were performed on a set of 100,000 subsampled SNPs.

**Table 2:** Difierent normalizations of the GTEx dataset.

| DataSet | Description |
| --- | --- |
| Remove PC | We keep removing first several (up to 300) principle components (PCs) until the network quality metrics (mean AUC and mean AUPR) no longer improve. |
| Remove tech | We remove the technical covariates by ridge regression with cross validation. |
| Remove tech + PC | We remove the technical covariates as above and subsequently remove residual PCs until the network performance metrics no longer improve. |
| DataRemix | DataRemix normalization is performed with $k$ ranging from 1 to 100. $p \in [-1, 1]$ and $\mu \in [0, 1]$ |
| HCP | HCP normalization is performed with following parameter settings. $k \in [1, 2, 3, 4, 5, 10, 20, 30, 40, 50, 60, 70, 80, 90, 100]$, $\lambda \in [1, 5, 10, 20]$, $\sigma_1 \in [1, 5, 10, 20]$ and $\sigma_2 \in [10, 20]$. We run grid search to pick up the best combination of parameters. |

### Parameter Optimization

The parameters *λ* = (*k, p, µ*) need to be optimized with respect to a particular biological objective. Grid search and random search (Bergstra & Bengio 2012) are among the most popular strategies, but these methods have low efficiency. Most of the search steps are wasted and the optimality of parameters is highly constrained by the step size and available computing power. In order to utilize the search history and keep a good balance between exploration and exploitation, we can formulate parameter search as a dual learning task.

We define a general performance measure *y* = *L*(*λ, 𝒟*), with *λ* representing the parameter tuple (*k, p, µ*), 𝒟 as the data, *L* as the evaluating process and *y* as the biological objective. Ideally we can determine the optimal point argmax_*λ*_ *L* easily by gradient descent based method, but usually *L* is derivative-free and it is also time intensive. Thus we introduce a surrogate model *f* (*λ*) which can directly predict *L*(*λ, 𝒟*) only given *λ*, and there are two conditions on *f* : argmax_*λ*_ *f* should be easy to solve and *f* should have enough capacity.

With these two properties, we can sequentially update *f* with (*λ*_*t*_, *y*_*t*_) and propose to evaluate *L* at *λ*_*t*+1_ = argmax_*λ*_ *f* in the next step. By gradually updating *f* with newly evaluated samples (*λ, y*), argmax_*λ*_ *f* approaches the true underlying optimal argmax_*λ*_ *L* as *f* can gradually fit to the underlying mapping function *L*. This provides a more efficient approach to explore the parameter space by exploiting the search history. In this work, we model *f* as a sample from a Gaussian Process with mean 0 and kernel *k*(*λ, λ*′), where *λ* = (*k, p, µ*)^*T*^. It is well known that the form of the kernel has considerable effect on performance. After experimentation we settled on the exponential kernel as the most suited for our application. The exponential kernel is defined as below (note the difference from the squared-exponential or RBF kernel).

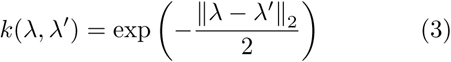

We observe *y*_*t*_ = *f* (*λ*_*t*_) + *ϵ* _*t*_, where *ϵ*_*t*_ ∼ *N* (0, *σ*^2^). For Bayesian optimization, one approach for picking the next point to sample is to utilize acquisition functions (Snoek *et al.*. 2012) which are defined such that high acquisitions correspond to potentially improved performance. An alternative approach is the Thompson Sampling approach (Basu & Ghosh 2017; Agrawal & Goyal 2013; Hernández-Lobato *et al.*. 2014). After we update the the posterior distribution *P* (*f*|*λ*_1:*t*_, *y*_1:*t*_), we draw one *sample f* from this posterior distribution as the optimization target to infer *λ*_*t*+1_. Theoretically it is guaranteed that *λ*_*t*_ converges to the optimal point gradually (Agrawal & Goyal 2013). With this theoretical guarantee, we focus on Thompson Sampling approach to optimize parameters for DataRemix.

#### Estimation of Hyper-Parameters

First we rely on the maximum likelihood estimation (MLE) to infer the variance of noise *σ*^2^ (Rasmussen 2004). Given the marginal likelihood defined by (4), it is easy to use any gradient descent method to determine the optimal *σ*^2^

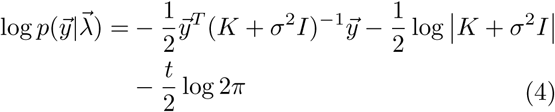

where 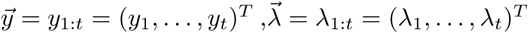 and *K* is the covariance matrix with each entry *K*_*ij*_ = *k*(*λ*_*i*_, *λ*_*j*_).

#### Sampling from the Posterior Distribution

Since Gaussian Process can be viewed as Bayesian linear regression with infinitely many basis functions *ϕ*_0_(*λ*), *ϕ*_1_(*λ*), … given a certain kernel (Rasmussen 2004), in order to construct an analytic formulation for the sample *f*, first we need to construct a certain set of basis functions Φ(*λ*) = (*ϕ*_0_(*λ*), *ϕ*_1_(*λ*), …), which is also defined as feature map of the given kernel. Then we can write the kernel *k*(*λ, λ*′) as the inner product Φ(*λ*)^*T*^ Φ(*λ*′).

Mercer’s theorem guarantees that we can express the kernels in terms of eigenvalues and eigenfunctions, but unfortunately there is no analytic solution given the exponential kernel we used. Instead we make use of the random Fourier features to construct an approximate feature map (Rahimi & Recht 2008). First we compute the Fourier transform *p* of the kernel (see Supplementary Methods for derivation).

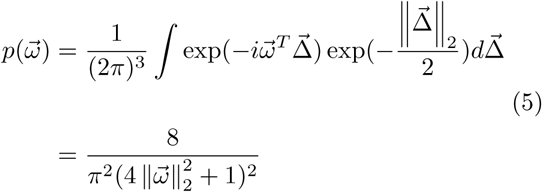

where 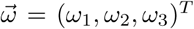 and 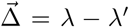. Then we draw *m*_*t*_ iid samples 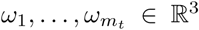 by rejection sampling with *p*(*ω*) as the probability distribution. Also we draw *m*_*t*_ iid samples 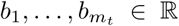 from the uniform distribution on [0, 2*π*]. Then the feature map is defined by the following equation.

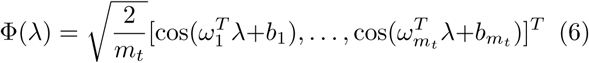

where the dimension *m*_*t*_ can be chosen to achieve the desired level of accuracy with respect to the difference between true kernel values *k*(*λ, λ*′) and the approximation Φ(*λ*)^*T*^ Φ(*λ*′).

#### Thompson Sampling

Any sample *f* from the Gaussian Process can be defined by *f* (*λ*) = Φ(*λ*)^*T*^ *θ*, where *θ* ∼ *N* (0, *I*) and Φ(*λ*)^*T*^ is defined by (6). In order to draw a posterior sample *f*, we just need to draw a random sample *θ* from the posterior distribution 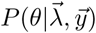.

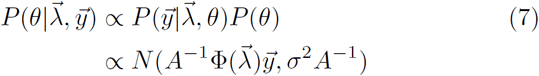

where 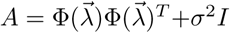 and 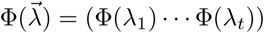. (see Supplemental Note for more details). The overall algorithm is summarized as the following pseudo code.

##### Algorithm 1 Thompson Sampling for Searching *λ*

Extra Parameters

*t*_*max*_: the maximum number of iteration steps

*ξ*: a pre-defined probability which ensures the search doesn’t get stuck in a local optimum

1. Get a short sequence 𝒟_1_ = (*λ, y*) as seeds by random search.
2. Draw *m*_*t*_ iid samples 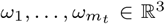 and *m*_*t*_ iid samples 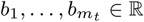 according to (5)
3. Iterate from *t* = 1 until *λ* converges or it reaches *t*_*max*_ (1) At step *t*, estimate the hyper-parameter *σ*^2^ given 𝒟_*t*_ according to (4) (2) Draw a sample *f* given 𝒟_*t*_ according to (7) with feature map determined by (6) (3)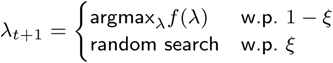 (4) Evaluate *y*_*t*+1_ given *λ*_*t*+1_ (5) 𝒟_*t*+1_ = 𝒟_*t*_ ⋃ (*λ*_*t*+1_, *y*_*t*+1_)

## Software availability

DataRemix is an R package which is freely available at GitHub (https://github.com/wgmao/DataRemix).

## Acknowledgements

This work was funded by the National Institutes of Health (R01 HG009299-01A1, U24 DK112331-01, U54 HG008540-03 and R03 MH109009-01A1 to M.C.). This study uses data from dbGaP (phs000486.v1). Funding support for the GAIN Major Depression: Stage 1 Genome-wide Association In Population Based Samples Study (parent studies: Netherlands Study of Depression and Anxiety (NESDA) and the Netherlands Twin Register (NTR)) was provided by the Netherlands Scientific Organization (904-61-090, 904-61-193, 480-04-004, 400-05-717, NWO Genomics, SPI 56-464-1419) the Centre for Neurogenomics and Cognitive Research (CNCR-VU); the European Union (EU/WLRT-2001-01254), ZonMW (geestkracht program, 10-000-1002), NIMH (RO1 MH059160) and matching funds from participating institutes in NESDA and NTR, and the genotyping of samples was provided through the Genetic Association Information Network (GAIN). The dataset(s) used for the analyses described in this manuscript were obtained from the database of Genotypes and Phenotypes (dbGaP) found at http://www.ncbi.nlm.nih.gov/gap through dbGaP accession number phs000486.v1.p1. Samples and associated phenotype data for the GAIN Major Depression: Stage 1 Genome-wide Association In Population Based Samples Study (PI: Dr. Patrick F. Sullivan, MD, University of North Carolina) were provided by Dr. Dorret I. Boomsma, PhD and Dr. Eco de Geus, PhD VU University Amsterdam (PIs NTR), Dr. Brenda W. Penninx, PhD, VU University Medical Center Amsterdam, Dr. Frans Zitman, MD PhD, Leiden University Medical Center, Leiden, and Dr. Willem Nolen, MD PhD, University Medical Center Groningen (PIs and site-PIs NESDA)

## Additional Files

Additional file 1 — SupplementaryFigures.pdf

Additional file 2 — SupplementaryMethods.pdf

