## Supplementary Methods for "DataRemix: a universal data transformation for optimal inference from gene expression datasets"

### Supplemental Methods

#### Fourier transform of the exponential kernel: equation(5)

The original paper which proposed to use Fourier features to construct an approximate feature map (Rahimi & Recht 2008) lists analytic formulations for three popular kernels, which are Gaussian kernel, Laplacian kernel and Cauchy kernel. Here we provide the detailed derivation for the Fourier transform of the exponential kernel in the three-dimensional space.

$$p(\omega) = \frac{1}{(2\pi)^3} \int \exp(-i\vec{\omega}^T \vec{\Delta}) \exp(-\frac{\left\|\vec{\Delta}\right\|_2}{2}) d\vec{\Delta}$$

First, we take the substitution $w = \|\vec{\omega}\|_2$ and $r = \left\|\vec{\Delta}\right\|_2$. We assume that $\vec{\omega}$ is parallel to the polar direction.

$$\begin{aligned}
p(\omega) &= \frac{1}{(2\pi)^3} \int \exp(-i\vec{\omega}^T \vec{\Delta}) \exp(-\frac{\left\|\vec{\Delta}\right\|_2}{2}) d\vec{\Delta} \\
&= \frac{1}{(2\pi)^3} \int_0^\infty \int_0^{2\pi} \int_0^\pi \exp(-iwr\cos\theta) \exp(-\frac{r}{2}) \cdot r^2 \sin\theta dr d\theta d\phi \\
&= \frac{1}{(2\pi)^3} \int_0^\infty r^2 dr \int_0^{2\pi} d\phi \int_0^\pi \exp(-iwr\cos\theta) \exp(-\frac{r}{2}) \cdot \sin\theta d\theta \\
&= \frac{1}{(2\pi)^2} \int_0^\infty r^2 dr \int_0^\pi \exp(-iwr\cos\theta) \exp(-\frac{r}{2}) \cdot \sin\theta d\theta
\end{aligned}$$

Here we make the substitution $z = \cos\theta$. Thus $\sin\theta d\theta = -dz$.

$$p(\omega) = \frac{1}{(2\pi)^2} \int_0^\infty r^2 dr \int_1^{-1} -\exp(-iwrz) \exp(-\frac{r}{2}) dz$$

Make another substitution $t = -iwrz$, where $dz = -\frac{1}{iwr}$

$$\begin{aligned}
p(\omega) &= \frac{1}{(2\pi)^2} \int_0^\infty r^2 dr \int_1^{-1} -\exp(-iwrz) \exp(-\frac{r}{2}) dz \\
&= \frac{1}{(2\pi)^2} \int_0^\infty r^2 dr \int_{-iwr}^{iwr} \exp(t) \exp(-\frac{r}{2}) \cdot \frac{1}{iwr} dt \\
&= \frac{1}{(2\pi)^2} \int_0^\infty r \exp(-\frac{r}{2}) dr \int_{-iwr}^{iwr} \exp(t) \cdot \frac{1}{iw} dt \\
&= \frac{1}{(2\pi)^2 \cdot iw} \int_0^\infty r \exp(-\frac{r}{2}) dr \int_{-iwr}^{iwr} \exp(t) dt \\
&= \frac{2}{(2\pi)^2 \cdot iw} \int_0^\infty r \exp(-\frac{r}{2}) dr \cdot i\sin(wr) \\
&= \frac{2}{(2\pi)^2 w} \int_0^\infty r \exp(-\frac{r}{2}) \sin(wr) dr
\end{aligned}$$

$$
\begin{aligned}
&= \frac{2}{(2\pi)^2 w} \cdot \frac{-4w\exp(-\frac{r}{2})(4w^2 r + r + 4)\cos(wr)}{(4w^2+1)^2}\Bigg|_0^\infty \\
&= \frac{2}{(2\pi)^2 w} \cdot \frac{16w}{(4w^2+1)^2} \\
&= \frac{8}{\pi^2(4w^2+1)^2} \\
&= \frac{8}{\pi^2(4\left\|\vec{\omega}\right\|_2^2+1)^2}
\end{aligned}
$$

#### Posterior sampling: equation(7)

The formula for sampling from the posterior can be intuitively understood in terms of the parallels with linear regression. Since in the feature space, any sample $f(\lambda)$ from Gaussian Process can be approximated by $\Phi(\lambda)^T\theta$, we can think of this as a simple linear regression: substitute $f(\lambda) = y - \epsilon, \epsilon \sim N(0,\sigma^2)$ and wish to solve $y = \Phi(\lambda)^T\theta + \epsilon, \epsilon \sim N(0,\sigma^2)$ for $\theta$. If $\theta$ comes with a Gaussian prior $N(0, I)$, then the posterior distribution of $\theta$ is $N(A^{-1}\Phi(\vec{\lambda})\vec{y}, \sigma^2 A^{-1})$, where $A = \Phi(\vec{\lambda})\Phi(\vec{\lambda})^T + \sigma^2 I$. The mean value $(\Phi(\vec{\lambda})\Phi(\vec{\lambda})^T + \sigma^2 I)^{-1}\Phi(\vec{\lambda})\vec{y}$ is the same as the ridge regression estimator or MAP (Maximum a posteriori estimation) estimator of $\theta$. For a full formal derivation see bellow:

$$
P(\theta|\vec{\lambda},\vec{y}) \propto P(\vec{y}|\vec{\lambda},\theta)P(\theta)
$$

where $P(\vec{y}|\vec{\lambda},\theta) \sim N(\Phi(\vec{\lambda})^T\theta, \sigma^2 I), P(\vec{\theta}) \sim N(0, I)$

$$
\begin{aligned}
P(\theta|\vec{\lambda},\vec{y}) &\propto \frac{1}{\sqrt{(2\pi)^t}\sigma^t}\exp\left(-\frac{1}{2}(\vec{y}-\Phi(\lambda)^T\theta)^T \cdot \sigma^{-2} I \cdot (\vec{y}-\phi(\lambda)^T\theta)\right) \cdot \frac{1}{\sqrt{(2\pi)^t}}\exp\left(-\frac{1}{2}\theta^T\theta\right) \\
&\propto \exp\left(-\frac{1}{2}(\vec{y}-\Phi(\lambda)^T\theta)^T \cdot \sigma^{-2} I \cdot (\vec{y}-\Phi(\lambda)^T\theta)\right) \cdot \exp\left(-\frac{1}{2}\theta^T\theta\right) \\
&= \exp\left(-\frac{1}{2\sigma^2}\left[\vec{y}^T\vec{y} - 2\vec{y}^T\Phi(\lambda)^T\theta + \theta^T\Phi(\lambda)\Phi(\lambda)^T\theta\right] - \frac{1}{2}\theta^T\theta\right) \\
&\propto \exp\left(-\frac{1}{2\sigma^2}\left[\vec{y}^T\vec{y} - 2\vec{y}^T\Phi(\lambda)^T\theta + \theta^T\Phi(\lambda)\Phi(\lambda)^T\theta\right] - \frac{1}{2}\theta^T\theta\right) \cdot \text{constant} \\
&= \exp\left(-\frac{1}{2\sigma^2}\left[\vec{y}^T\vec{y} - 2\vec{y}^T\Phi(\lambda)^T\theta + \theta^T\Phi(\lambda)\Phi(\lambda)^T\theta\right] - \frac{1}{2}\theta^T\theta\right) \\
&\cdot \exp\left(-\frac{1}{2\sigma^2}\vec{y}^T\left[\frac{1}{\sigma^2}\Phi(\lambda)^T(\frac{1}{\sigma^2}\Phi(\lambda)\Phi(\lambda)^T - I)^{-1}\Phi(\lambda) - I\right]\vec{y}\right) \\
&= \exp\left(-\frac{1}{2}(\theta-\hat{\theta})^T(\frac{1}{\sigma^2}\Phi(\lambda)\Phi(\lambda)^T + I)(\theta-\hat{\theta})\right)
\end{aligned}
$$

where $\hat{\theta} = \frac{1}{\sigma^2}(\frac{1}{\sigma^2}\Phi(\lambda)\Phi(\lambda)^T + I)^{-1}\Phi(\lambda)\vec{y}$. Thus,

$$
\begin{aligned}
P(\theta|\vec{\lambda},\vec{y}) &\propto \exp\left(-\frac{1}{2}(\theta-\hat{\theta})^T B(\theta-\hat{\theta})\right) \\
&\propto N(\frac{1}{\sigma^2}B^{-1}\Phi(\vec{\lambda})\vec{y}, B^{-1})
\end{aligned}
$$

where $B = \frac{1}{\sigma^2}\Phi(\vec{\lambda})\Phi(\vec{\lambda})^T + I$. Here, $B$ is a simple transformation of the previously defined quantity $A = \Phi(\vec{\lambda})\Phi(\vec{\lambda})^T + \sigma^2 I$ with $B = \frac{1}{\sigma^2}A$. Equivalently, we get,

$$P(\theta|\vec{\lambda}, \vec{y}) \propto N(A^{-1}\Phi(\vec{\lambda})\vec{y}, \sigma^2 A^{-1})$$

where $A = \Phi(\vec{\lambda})\Phi(\vec{\lambda})^T + \sigma^2 I$.
