## Supplementary Figures for "DataRemix: a universal data transformation for optimal inference from gene expression datasets"

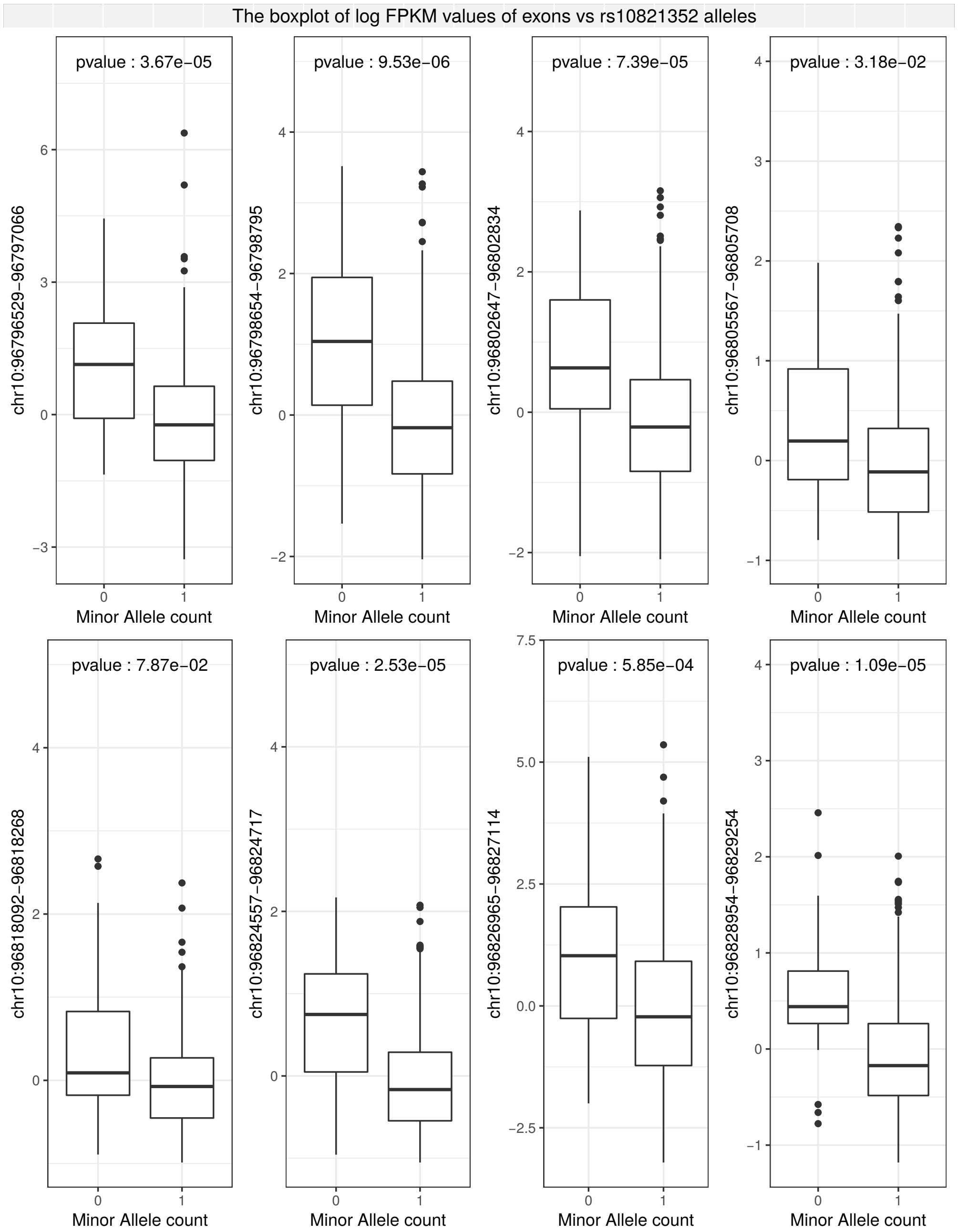


Figure S1: The effect of rs10821352 on CYP2C8 expression is consistent across exons. We used raw FPKM data to quantify the exon-level trans-eQTL effects. The effect is consistent across exons further confirming that it is unlikely to be due to homolog mismapping and other technical artifacts.

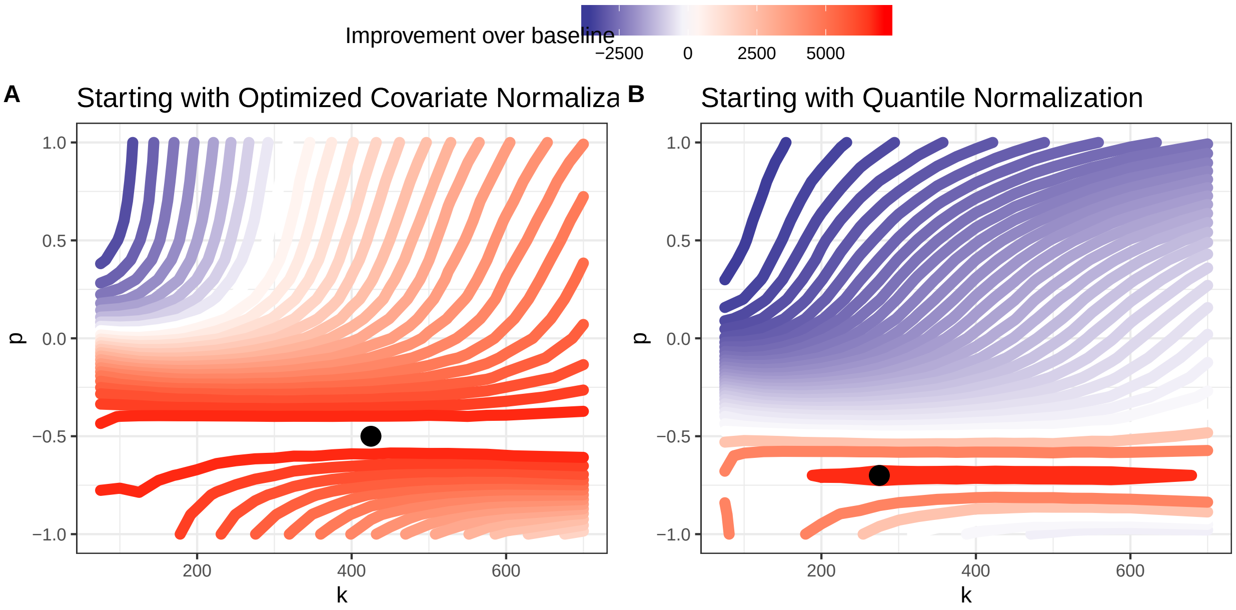


Figure S2: Contour plot representing the effects of the $k$ and $p$ parameters on the performance of DataRemix regarding cis-eQTL discovery on training set. The $\mu$ parameter is fixed at 0.01. Red contours represent parameter combinations that increase the number cis-eQTLs beyond what can be achieved using the $D_{HCP-cis}$ dataset. Panel A shows the results starting with $D_{cis-optim}$ while $D_{QN}$ is used for panel B. Improvement can be achieved starting with either datasets. We note that the optimal $p$ parameter is negative (though slightly different) for both datasets.

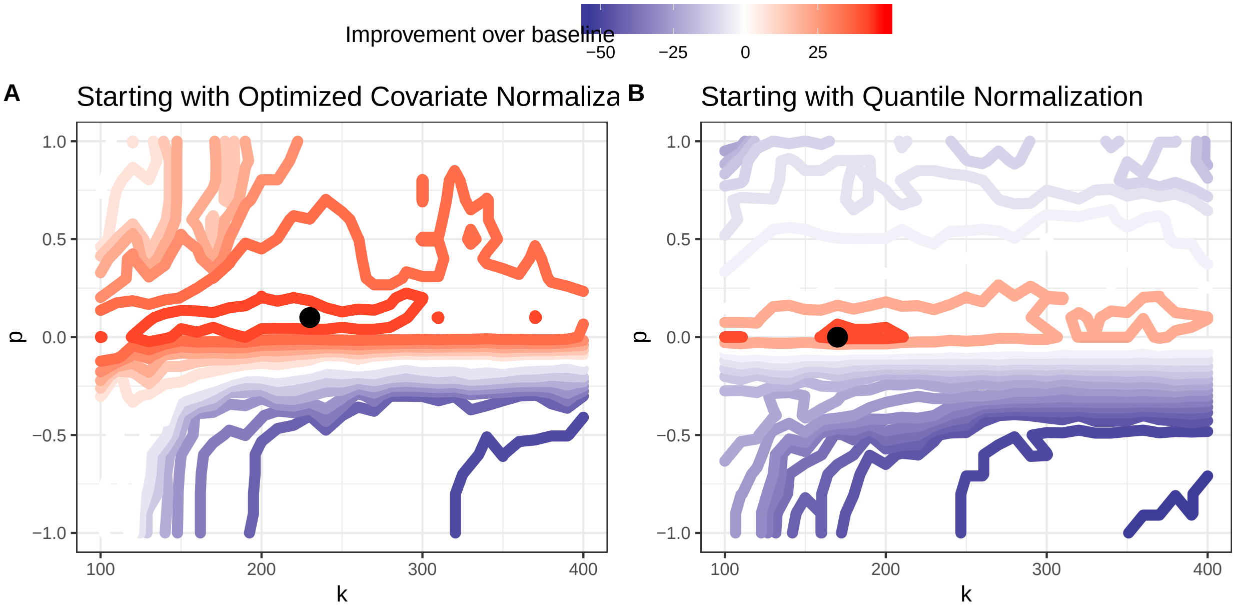


Figure S3: Contour plot representing the effects of the $k$ and $p$ parameters on the performance of DataRemix regarding trans-eQTL discovery on training set. The $\mu$ parameter is fixed at 0.01. Red contours represent parameter combinations that increase the number trans-eQTLs beyond what can be achieved using the $D_{HCP-trans}$ dataset. Panel A shows the results starting with $D_{HCP-trans}$ while $D_{QN}$ is used for panel B. Improvement can be achieved starting with either datasets. We find that the region of improved performance is smaller than for cis-eQTLs and is particularly concentrated when starting with the $D_{QN}$ (panel B) dataset.
